# Guadalupe fur seal alopecia: a metabolic syndrome associated to climatic anomalies?

**DOI:** 10.1101/2023.04.02.535308

**Authors:** Ariadna A. Guzmán-Solís, Fernando R. Elorriaga-Verplancken, Karina A. Acevedo-Whitehouse

## Abstract

Alopecia is characterized by the thinning and loss of hair or fur, and has been observed in different marine carnivores. Until recently, there were no records of alopecia in marine mammals from the Northeast Pacific Ocean. However, sightings of juvenile male Guadalupe fur seals with patchy alopecia in the ventral body have been reported in the San Benito Archipelago. This is concerning given that the species is considered endangered under Mexican law and has only started to recover from near extinction less than a century ago. We captured 13 fur seals at San Benito Archipelago during the summer of 2017 and 2018 and collected fur samples for common dermatological analysis, as well as scanning electron microscopy and X-ray spectroscopy. We found marked damage to the structure of guard fur, with loss of integrity, medullary damage and perforations. Damage to secondary underfur hair was less common. We found no evidence of dermatophyte microorganisms or ectoparasites commonly associated with alopecia. We suggest that the damage is caused by metabolic alterations secondary to nutritional stress that is consequential to high sea surface temperatures that alter prey availability. This is the first study of alopecia in an otariid pinniped of the Northeast Pacific Ocean.

## 1. Introduction

Alopecia syndrome has been reported for a number of marine mammal species, including polar bears, *Ursus maritimus* [1], grey seals, *Halichoerus grypus* [2], and a number of fur seals, such as the Australian fur seal, *Arctocephalus pusillus doriferus* [3], and the Subantarctic fur seal, *A. tropicalis* [4]. This syndrome, first reported in 2015 for Guadalupe fur seals, *A. townsendi*, is characterized by the thinning and subsequent loss of hair [5].

Being a syndrome, alopecia does not have a single aetiology. However, various factors have been correlatively associated with this condition. These include the presence of parasites, such as *Anoplura* lice and *Demodex zalophi* mites, which can lead to overkeratinization, destroy the epithelium and damage the hair follicles directly and can lead to anaemia [6]. Fungi [4,7], bacteria, and viruses could also lead to alopecia if they cause dermal lesions, abscesses or vesicles [2,8]. However, other factors can also play a contributory role. These include PCB pesticides [9], endocrine alterations [10], nutritional deficiencies that stunt growth and integrity of hair and skin structures [11–14], toxins [15], and immune dysfunction [16]. Whether these factors need to be present in all cases of alopecic syndrome is unknown.

Cases of alopecia began to be observed in Guadalupe fur seals during the atypically high sea surface temperature (SST) event of 2015 [5], coinciding with the start of the unusual mortality event recorded for the species and that lasted through 2021 [17]. Specifically, stranded pups and juveniles with alopecia were registered along the Oregon coastline [5], and there were sightings of juveniles with alopecia in the San Benito Archipelago, SBA [18]. This is concerning, given that Guadalupe fur seals were considered commercially extinct less than 100 years ago, and the species is listed as endangered and considered a priority species for conservation by the Mexican government [19], and while the population has increased in size and in distribution [20], the majority of its population and breeding colonies are located on a single island, Guadalupe Island, in the Mexican northern Pacific [21], with non-reproductive recolonization of other areas, such as SBA [22]. Furthermore, the species is susceptible to climatic anomalies [17,22,23], as it is dependent on its double layer of fur for thermoregulation [5], so conditions that affect its pelage, potentially causing alopecia, could be important in relation to their fitness, body condition, and health, as has been reported for other species, such as the polar bear [1].

Here, we use microscopic techniques to examine fur samples collected from Guadalupe fur seals with alopecia, to examine whether there is evidence of structural alterations in their fur. To the best of our knowledge, this is the first study of alopecia in this species.

## 2. Materials and methods

### (a) Capture and collection of hair

During July of 2017 and 2018, 13 juvenile male Guadalupe fur seals (*Arctocephalus townsendii*) were captured during routine monitoring of the species at SBA (28° 18 ‘ N, 115° 32 ‘ W) in the western region of the Baja California Peninsula. Individuals were captured using hoop nets and were physically restrained during the sampling procedure.

Fur (a 1.5 × 1.5 cm patch) was collected from areas that showed alopecia, and always in the ventral surface of the chest. We used sterile scissors for collection and kept the samples in small paper envelopes. Nitrile examination gloves were used throughout the procedure. For individuals that did not show detectable alopecia, fur samples were collected from the same body region and using the same methodology.

### (b) KOH Staining and UV illumination

A subset of each sample (at least, ten guard hairs with several secondary hairs per individual) were deposited on clean glass microscope slides and drops of KOH solution (containing 20% KOH and 40% DMSO in sterile deionized water) were added until the entire sample was covered. The slide was examined under an optical microscope at 10X, 40X, and 100X magnification. In addition, the hair strands were placed in Petri dishes to be examined by long-wavelength UV light under the stereoscopic microscope.

### (C) Scanning Electron Microscopy (SEM) and X-ray spectroscopy (EDS)

To determine hair shaft morphology, scanning electron microscopy (SEM) images of were taken of four to six guard hairs with several secondary hairs from each sample using a EVO50 (Carl Zeiss AG) SEM with variable pressure. Acceleration voltage was 20 kV, magnification at 40X and 100X. In order to identify the chemical composition of fur, we performed X-ray spectroscopy (EDS) on every sample that showed damage under SEM, using SE1, BSD, and EDX detectors. All analyses were conducted at the Laboratory of Microscopy of the School of Natural Sciences of the Autonomous University of Queretaro (Mexico). All samples were handled with sterile gloves to avoid any contamination.

## 3. Results

We observed internal structural differences between the fur seal coarse guard fur collected from zones with alopecic lesions and from zones with normal fur. Half of the examined guard fur analysed had damage regardless of alopecia being evident. Namely, normal guard fur showed cuticular scales well preserved on the periphery and an organized medullar column (Fig. 1A), while those from alopecic zones had marked medullary disorganization and areas of breakage or splintering of the hair (Fig. 1B). There was no other observable difference in the guard fur under light microscopy. Mostly, underfur showed no observable anomalies, with few secondary hairs revealing loss of structure and scales (Fig. 2).

**Figure 1.**
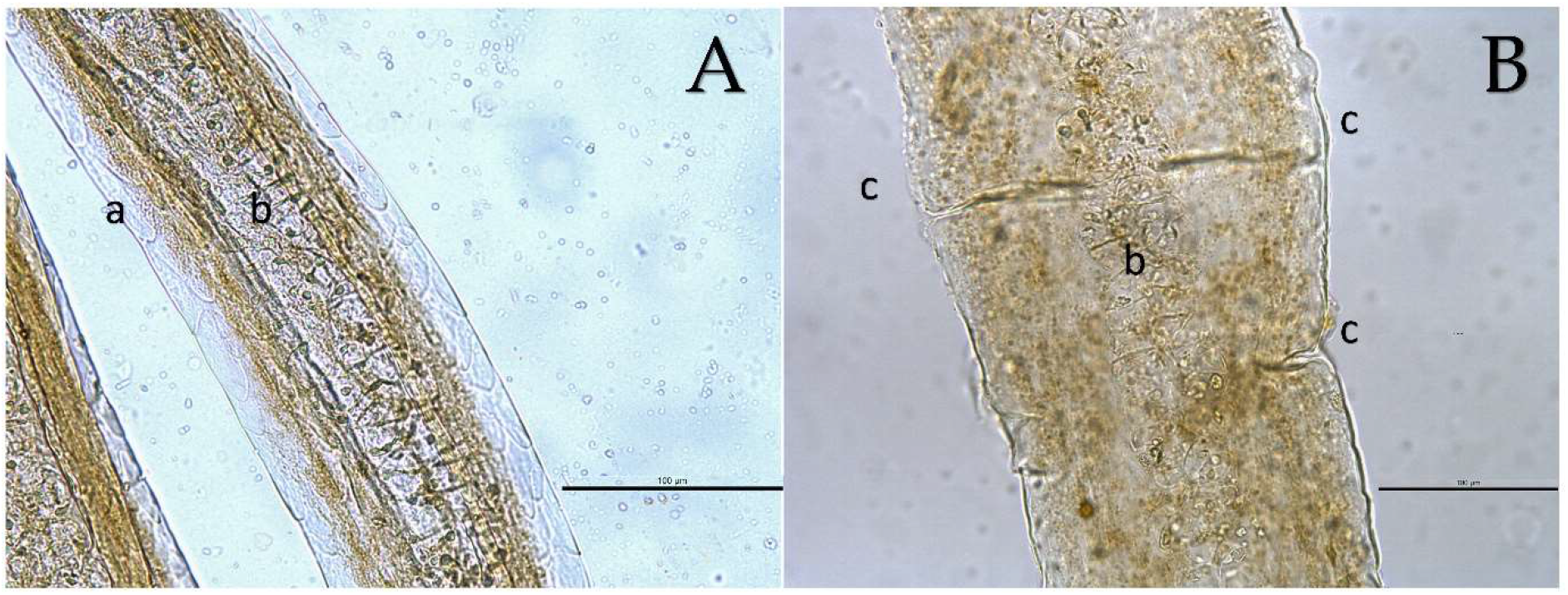
Microscopy of Guadalupe fur seal guard hair with potassium hydroxide (KOH). A) Guard hair showing preservation of cuticular scales at the periphery (a) and a well-organized medullar column (b); B) Guard hair showing remains of the medullary column without organization (b) and areas of splintering (c). Scale bar: 100 μm. Magnification 20X.

**Figure 2.**
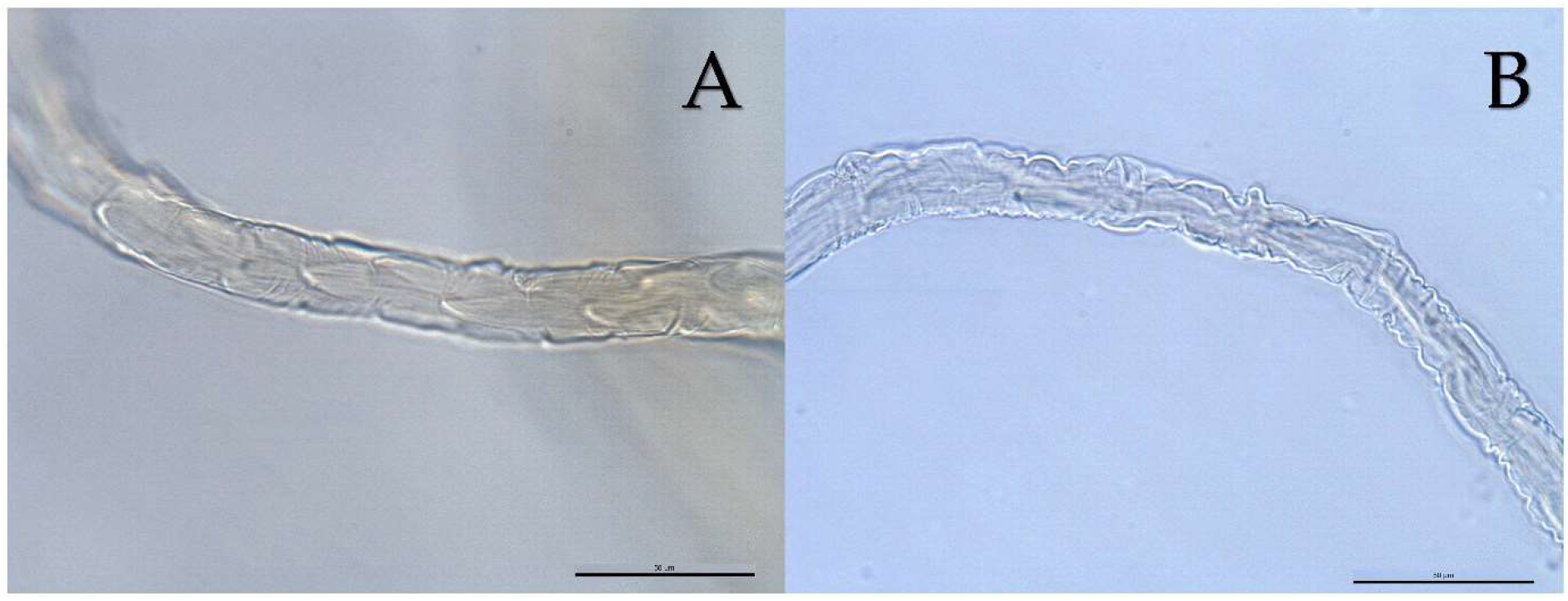
Microscopy of Guadalupe fur seal secondary hair with potassium hydroxide (KOH). A) cuticular structure showing preserved toothed scales B) secondary hair without scales, with break and irregular edges at the margin. Scale bar: 50 μm. Magnification 40X.

There was no evidence of fungal spores, septa, asci, yeast, or germ cells. We did not find parasites nor parasite eggs, neither in the guard hairs or the secondary fur. However, most of the guard fur (Fig. 3A) and some underfur (secondary) hairs (Fig. 3B) had unexpected structures accumulated along the shaft. UV illumination did not reveal any sign of fluorescence as would be expected by the presence of fungi or dermatophyte bacteria associated with fur loss.

**Figure 3.**
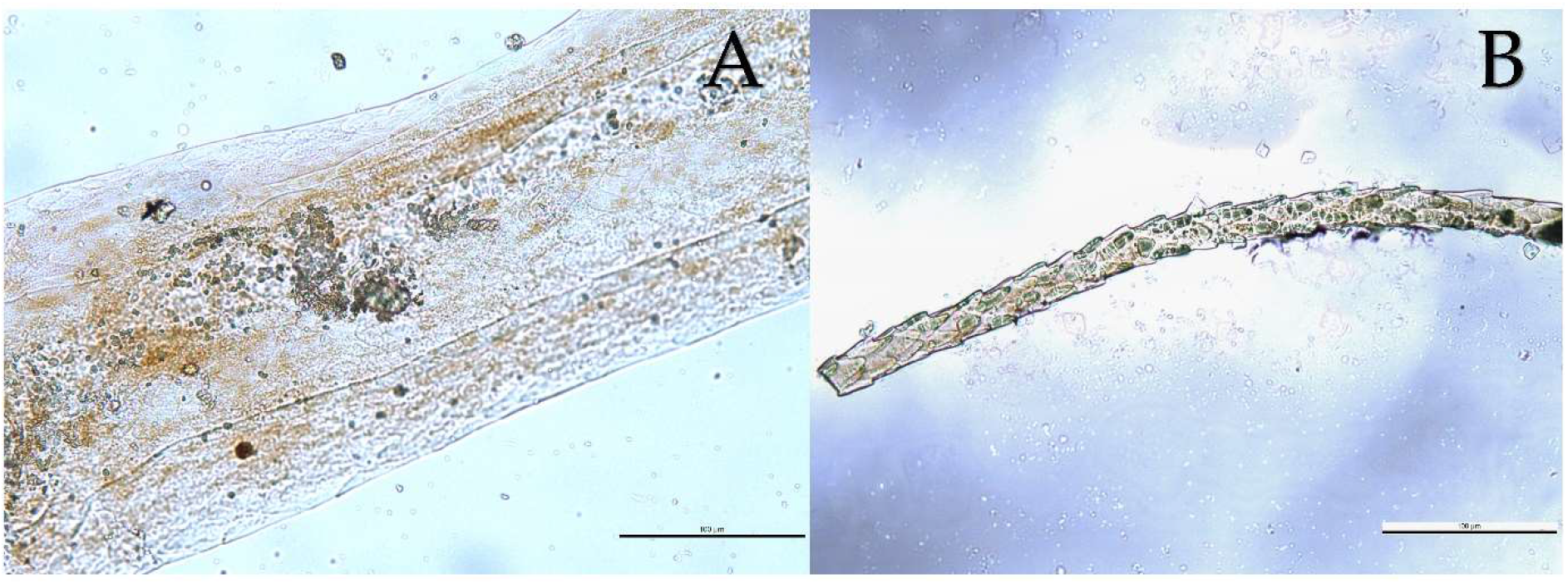
Microscopy of Guadalupe fur seal hair with potassium hydroxide (KOH).A) Guard hair and B) Secondary hair, both with accumulation of objects that do not reflect expected hair structures. Scale bar: 100 μm. Magnification 20X.

Scanning electron microscopy showed guard hairs with a conserved and well-organized cuticular structure of juxtaposed scales of the coronal dentate type (Fig. 4A), and guard hairs with structural damage (Fig. 4B). Interestingly, all fur seals had at least one hair with structural damage, regardless of showing evidence of alopecia. We found no evidence of ultrastructural damage in any of the underfur (secondary hair) samples analysed (Fig. 5).

**Figure 4.**
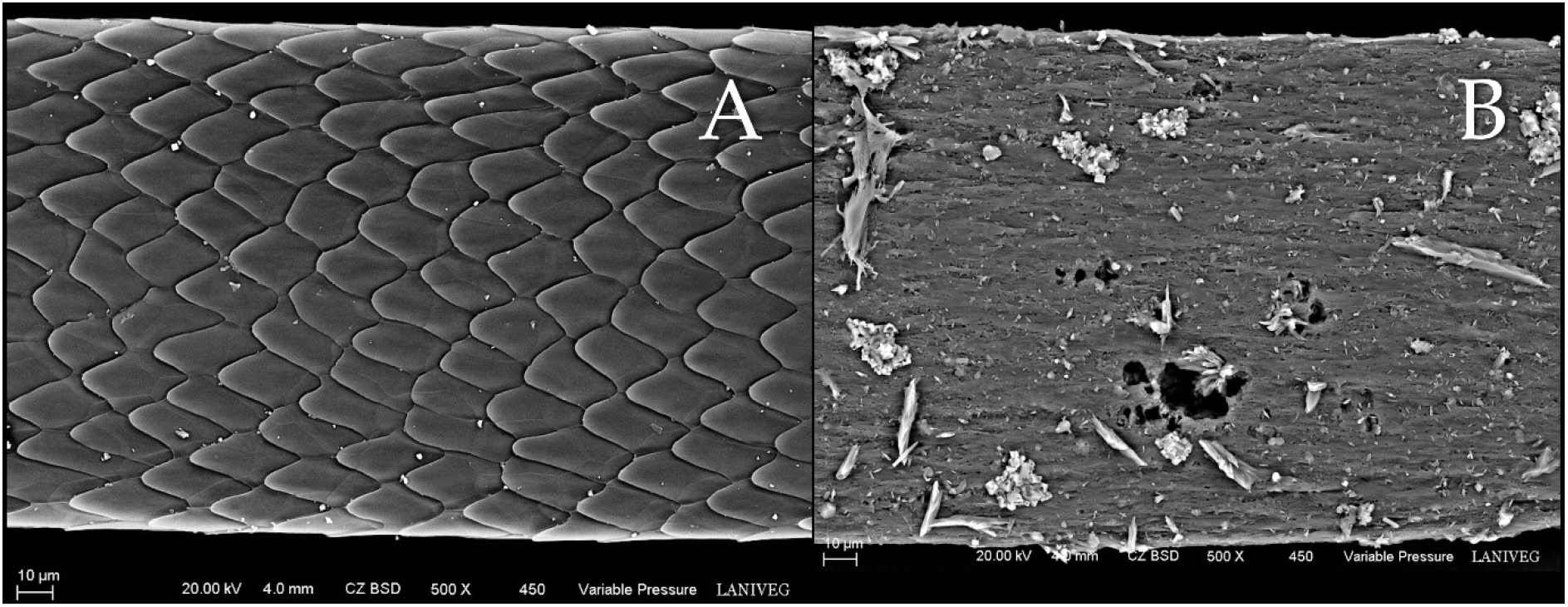
Scanning electron micrographs of A) Well-conserved cuticular scales in Guadalupe fur seal guard hair. B) Marked loss of cuticular scales, perforations, incrustations, and loss of margin in guard hair. Scale bar: 10 μm. Magnification 500X.

**Figure 5.**
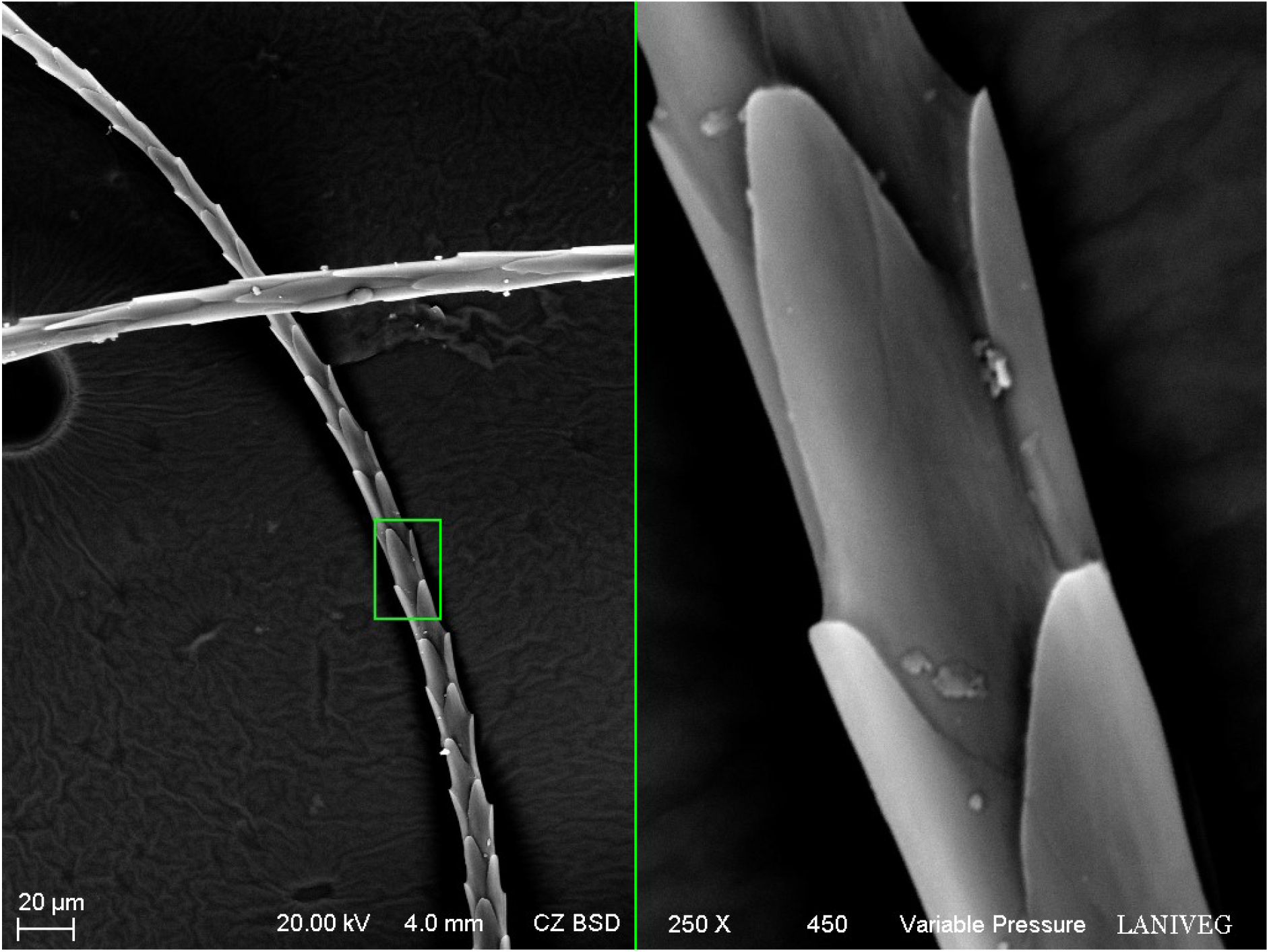
Scanning electron micrographs of the ultrastructure of a secondary hair with long dentate scales. Right panel shows a magnified (250X) image of the dentate scales. Scale bar: 20 μm.

Guadalupe fur seal guard fur showed different cuticular ultrastructure (Fig. 6), with some hair samples revealing well-conserved structures and uniform scales (Fig. 6A), others showing cuticular scales with no tip and diffuse organization in some parts (Fig. 6B), hairs with no scales, perforations and small loss of margin (Fig. 6C), and even fur with absence of cuticular scales, perforations, incrustations and marked loss of margins that fractured the hair transversally (Fig.6D and Fig. 7).

**Figure 6.**
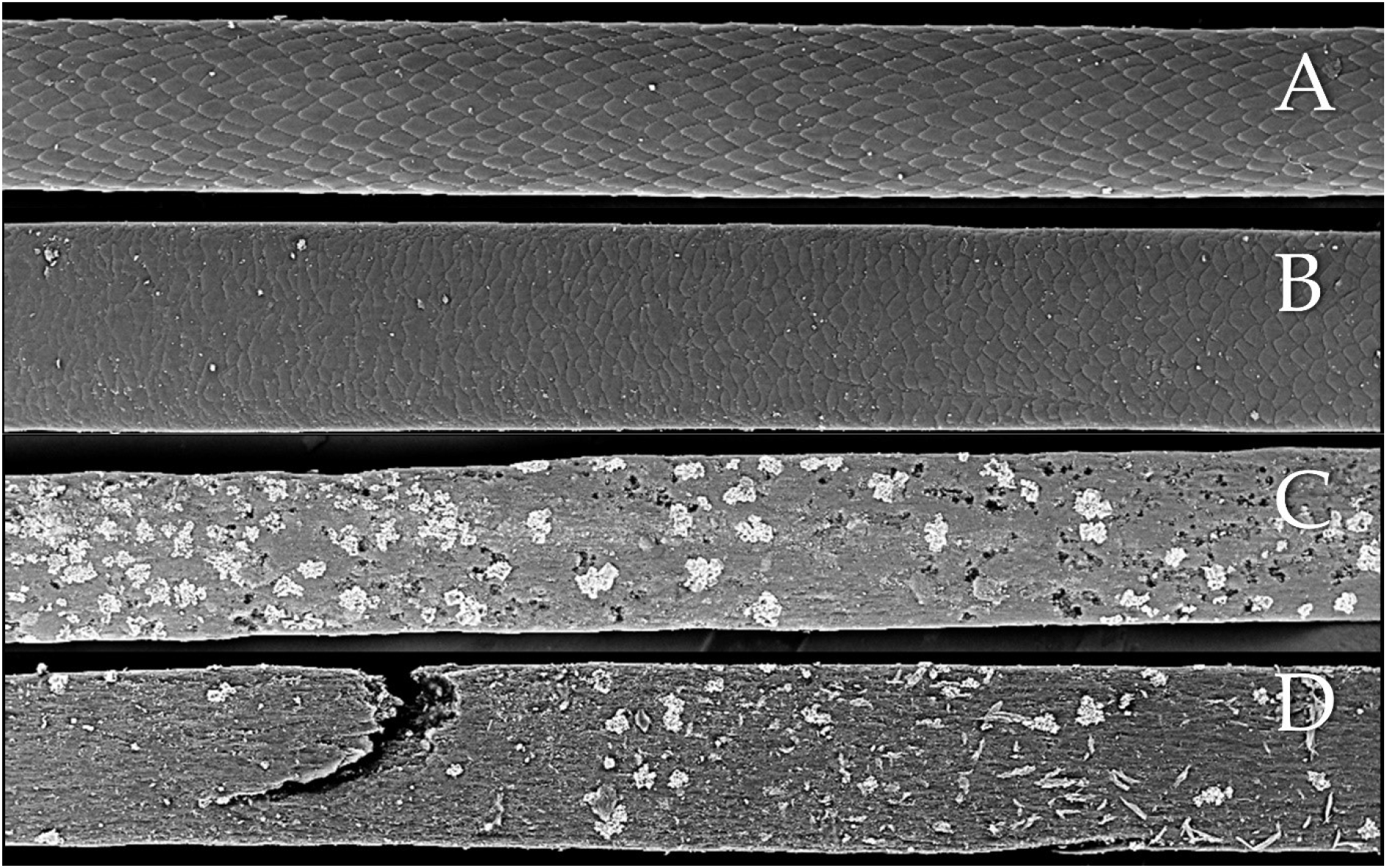
Ultrastructural microscopy of Guadalupe fur seal guard hairs showed different levels of loss of structural integrity. A) Guard hair with uniform scales and normal structure. B) Guard hair with mild loss of definition in scales. C) Guard hair with some perforations and encrusted foreign material. D) Severe damage to guard hair, with many perforations, encrusted material and ruptured shaft.

**Figure 7.**
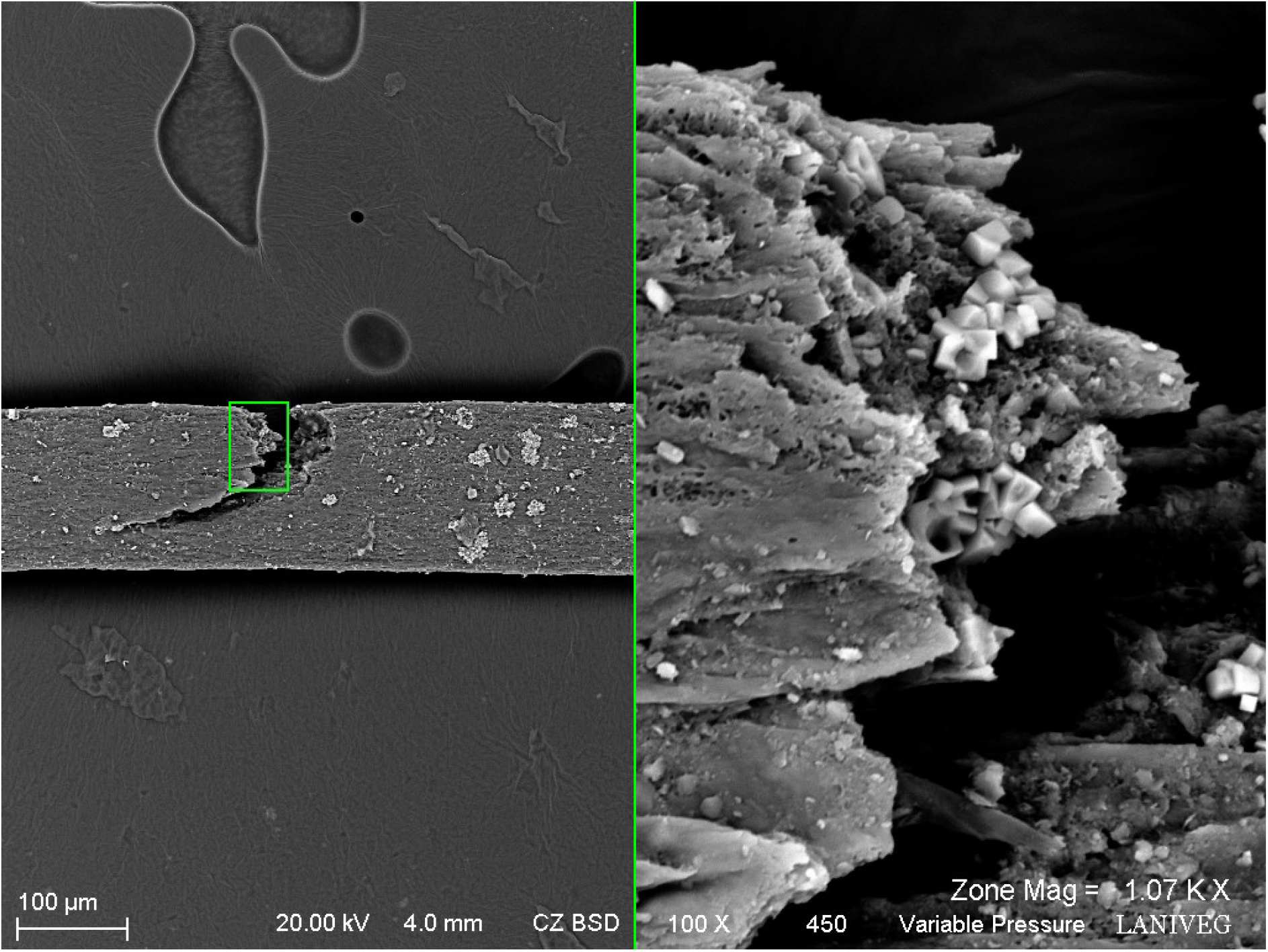
Scanning electron microscopy images of guard hair. Left panel shows the transversal fracture of the hair. Right panel shows a magnified (100X) image the fractured area. Scale bar: 100 μm.

Some of the guard fur with damage showed perforations in the cuticle which were covered with encrusted crystal-like material that did not show similarities to fungal, parasitic or bacterial structures (Fig. 8). X-ray dispersion analysis (EDS) revealed the particles to be formed by sodium (Na), calcium (Ca) and chloride (Cl) (Fig. 9).

**Figure 8.**
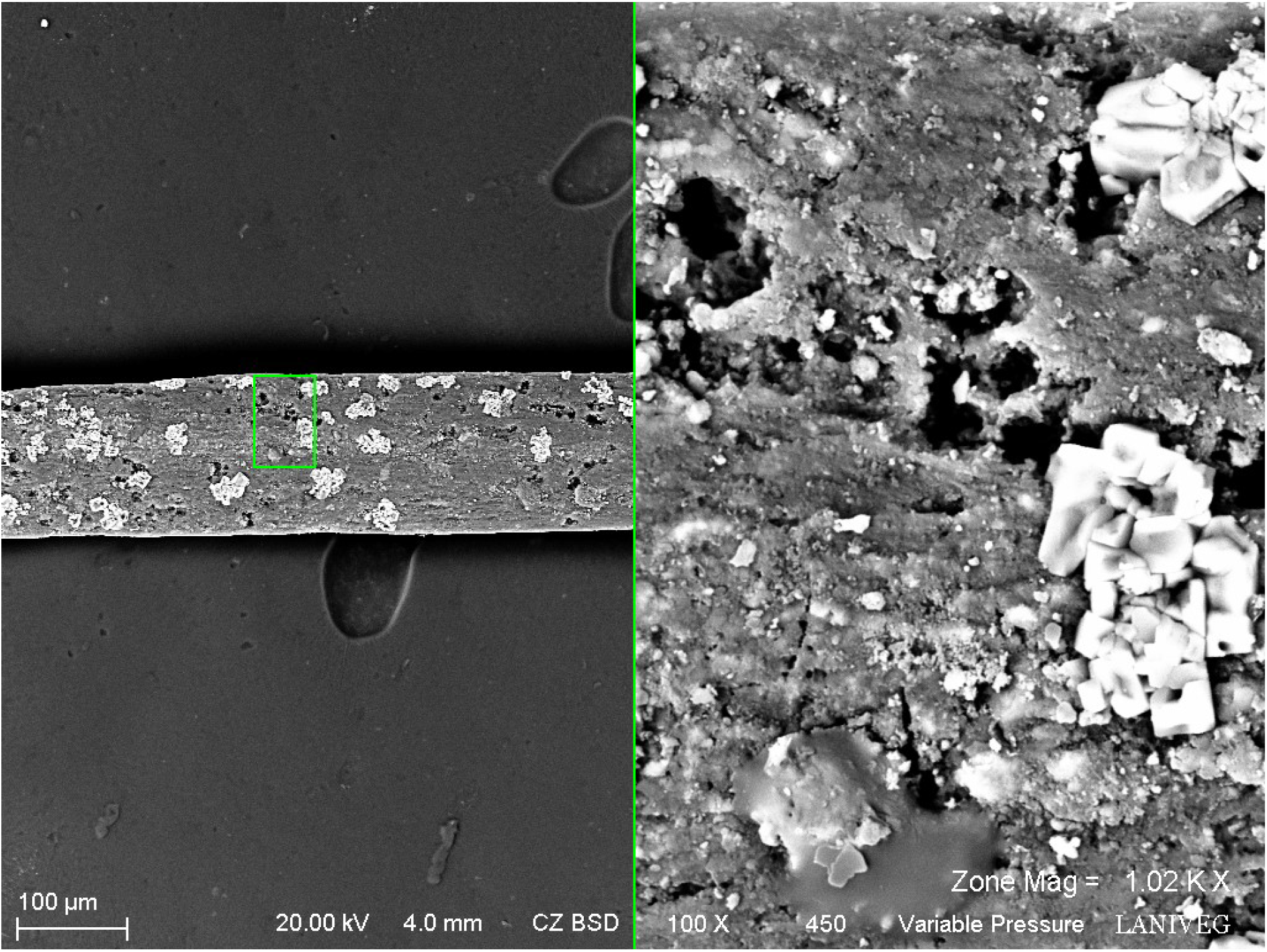
Scanning electron micrograph of perforations and encrusted particles found in Guadalupe fur seal guard hairs. Right panel shows a magnified (100X) image of the encrusted particles. Scale bar: 100 μm.

**Figure 9.**
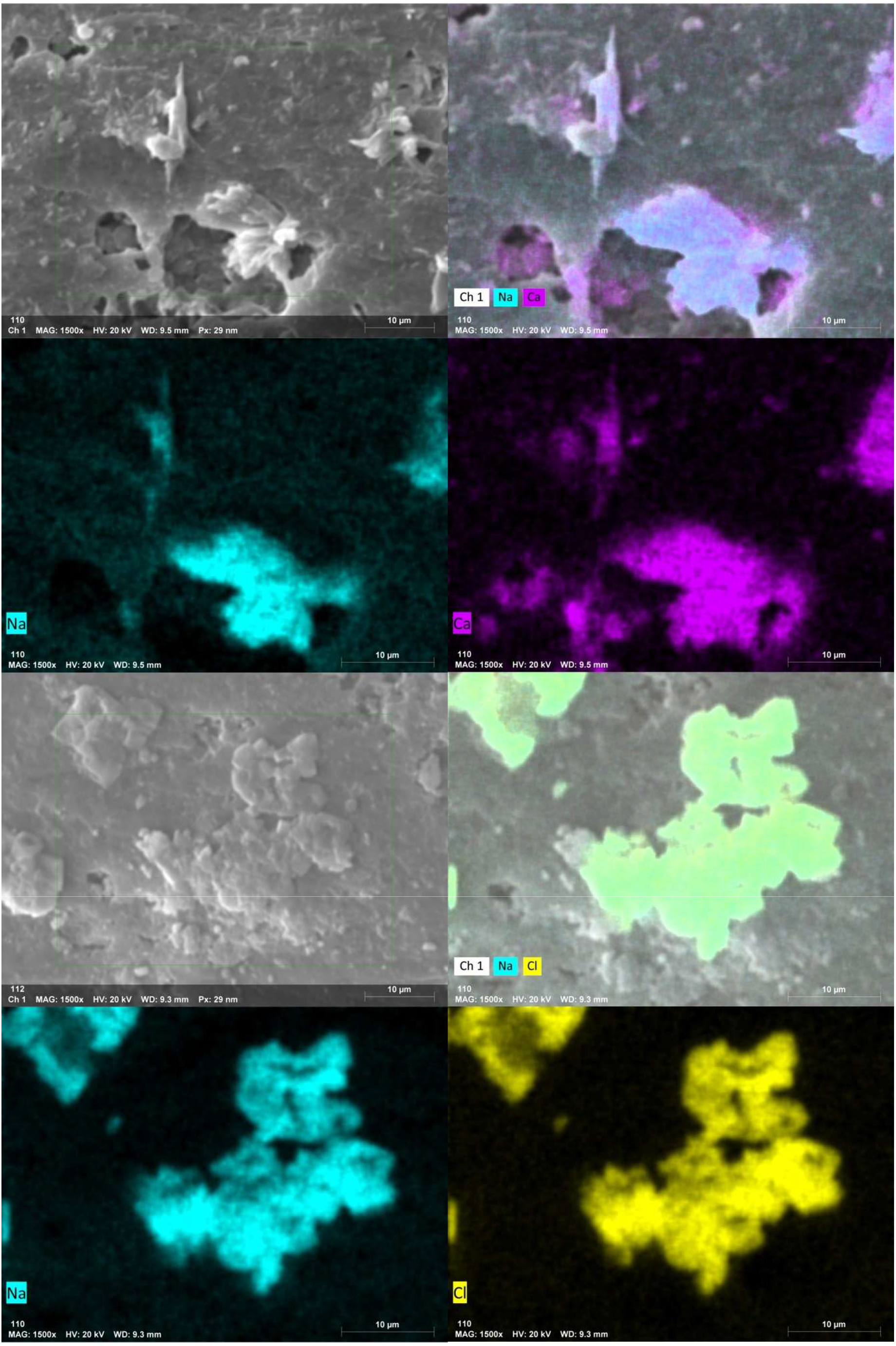
EDS analysis of the elemental composition of the encrusted material found with the perforations in the Guadalupe fur seal guard hair. Particle composition was sodium (Na), calcium (Ca) and chloride (Cl).

## 4. Discussion

The loss of pelage or fur damage can be a threat to fur seals, as their blubber thickness is modest compared to that of other species [24], making them dependent on their pelage as their main thermoregulatory mechanism. Fur seals have a thick double layer of fur that bidirectionally controls heat conduction and convection between the body and the environment [25,26]. When air is trapped within the fur layers, it forms a *shield* between the highly-irrigated skin and the environment, as well as aiding flotation [27]. This is why the loss of integrity of a fur seal ‘s pelage is not a trivial matter. When it is compromised, the underlying skin is exposed to the environment, decreasing the ability to thermoregulate, and increasing the exposure to environmental agents, including ultraviolet radiation [28] that can exert local and systemic damage [29]. Alopecia has been observed for other fur seal species: however the first cases of alopecia in Guadalupe fur seals were noted less than a decade ago during an unusual mortality event (UME) [5], which coincided with a period of anomalously high sea surface temperatures (SST), termed ‘The Blob ‘ or North Pacific heat wave, which started in 2014 and was worsened by the 2015 El Niño event [30], yielding increases in SST of more than 2.5°C [31]. We have studied the fur of juvenile males with evident alopecia and our study, to the best of our knowledge, is the first report of its kind for this age group, and the first report of alopecia in Guadalupe fur seals in the San Benito Archipelago, which is significant within the recovery process of the species as this is a recolonization site that harbours the second largest colony of Guadalupe fur seals (up to 3,700 individuals) after Guadalupe Island.

The guard hairs of the fur seals that we examined showed structural alterations and ultrastructural evidence of fragility and erosion. Hair is formed by fibrous proteins, mainly alpha keratin, whose amino acids include cysteine monomers [32]. Where their sulfhydryl groups join, disulphide bridges are formed, bestowing hair its shape and resistance in such a way that if cysteine is reduced, the disulphide bridges are broken and hair becomes brittle and changes it structure [33]. Other factors that can affect hair structure is exposure to high temperatures [34,35] as they modify the helical components of keratin [36], as well as direct contact or oral exposure to certain chemicals, such as heavy metals, selenium and boric acid, among others well known to cause alopecia in humans [37]. We propose that the structural damage to the fur of Guadalupe fur seals has a metabolic origin, secondary to climatic stressors, rather than being due to direct thermal stress to the fur. This is because thermal damage to hair occurs with exposure to temperatures as high as 235°C [38].

To explain why damage to the fur of Guadalupe fur seals could have a metabolic origin, we must focus our attention on cysteine and keratin, structural proteins that five resistance and stability to the fur coat. In particular, the synthesis of cysteine could be key to alopecia. This is because it depends on the availability of methionine, an essential amino acid that is obtained in the diet, mainly from items that have a high protein content and good quality [39]. The Guadalupe fur seal is vulnerable to the climatic anomalies that have been registered for its range in recent years [23,40]. If the Guadalupe fur seal ‘s preferred prey is negatively impacted by high SSTs, and the species switches to other prey of lower nutritional value, synthesis of cysteine could become compromised, with physical impacts, such as the observed alopecia. Interestingly, structural irregularities in guard hairs of fur seals whose fur appeared to be normal (i.e. they did not have evident alopecia) were also present. This finding leads us to hypothesize that the initial stages of the process could be asymptomatic and will only culminate with alopecia if the contributory factors persist. We propose that, once the normally resistant structure of guard hair is impacted following alteration of normal biochemical pathways, physical damage to the hair shaft is easier to occur, as seen by the perforations we observed, that were associated to the presence of Na, Ca, and Cl, the main components of sea salt. Salt granules could become trapped in the perforations, further increasing damage to fur, and leading to their breakage, as we observed in some samples.

The majority of damage observed occurred in the guard fur (primary hair), with limited signs of damage to the underfur (secondary hair). There are structural differences in the thickness, resistance and composition of guard and secondary fur of otariids. Specifically, secondary fur lacks a medullary composition, making guard hair more resistant [41]. Fur seal guard hair is coarse and particularly resistant to physical abrasion due to its ultrastructure, which renders them nearly unscathed even when undergoing digestion or carcass decomposition [42]. We propose that the limited damage of secondary hair was because the guard hair, even though damaged, kept offering protection to the underfur and only in a few individuals damage was so severe that the secondary hair began to exhibit signs of damage. It is possible that when the damage is extreme, both the guard and the secondary hair would be lost due to fragility and abrasion.

The observed damage to the medullary column and cuticular scales warrants careful discussion due to its potential impact on fur seals. The medullary column is a structural part of the hair and could also be affected by the availability of essential amino acids that are necessary for the synthesis and turnover of cysteine and keratin. Changes in the medullary structure of the fur impacts the ability to thermoregulate and to block solar radiation from reaching the skin, thus avoiding damaging radiation and dissipating heat [28]. These dissipation structures were damaged in the areas that exhibited alopecia. Given that alopecic patches were observed in the ventral chest and abdominal area, both highly vascularized and with arteriovenous anastomosis system (AVA), in contrast to other parts of the body of otariid pinnipeds [43]. Thus, fur medullary damage is likely to affect the species ‘ main mechanism of radiation and temperature dissipation and lead to overheating [28,44]. In addition, cuticular scales, also observed to be affected, are important for insulation of fur seals. This is because their elongation, smoothness and cuticular waxes are critical for keeping the fur coat waterproof, allowing insulation during and after immersions [45].

If our explanation of the observed structural abnormalities is indeed explained by metabolic alterations secondary to climate-related ecological disturbances, the consequences are likely to be far deeper than fur. On one hand, methionine, a precursor for cysteine synthesis, is a promoter of iron-sulphide groups and of disulphide bridges; these, in turn, are necessary for various physiological processes, including the joining of immunoglobulin light and heavy chains [33]. It is known that the effector activities of immunoglobulins are inhibited by the reduction of disulphide bridges. As well as the binding capacity of proteins in the complement system pathway that make the association of the antibody-antigen complex. Both cysteine and disulphide bridges also play a role in the structural stability of hormones such as insulin [46]. Also, suboptimal nutrition, possibly driven by changes in prey availability, will lower the amounts of resources available to an individual. Loss of fur will demand more energetic resources to maintain thermal homeostasis [47], in turn reducing the available resources for other costly physiological processes [48,49]. This has already been shown for other pinniped species (namely, the California sea lion, *Zalophus californianus*) that inhabit the same region, where individuals impacted by temperature-related dietary shifts had reduced inflammatory responses and antibody levels [50] and red blood cell membrane alterations [51].

Some studies conducted in other pinniped species have found parasites associated with alopecia [4,6]. However, these were done on captive individuals, and it is likely that their presence, and subsequent damage to the fur, was secondary to stress, as is known to occur in captivity for other species [52]. Free-living pinnipeds commonly harbour ectoparasites in their fur and skin, but these organisms do not tend to lead to disease unless the host ‘s immune response is suboptimal [53], as is well known for other mammals [54,55]. We found no evidence of lice, mites or other arthropod ectoparasites in the skin patch from where the fur samples were collected, nor did we find evidence of any ectoparasite attached to the collected fur. Beyond arthropod ectoparasites, dermatophyte fungi and bacteria can colonize tissues and structures that have keratin, such as hair and skin, leading to hypersensitivity reactions, inflammation, and subsequent dermal lesions and hair loss [7]. Using well-established microscopic and staining techniques to detect dermatophyte microorganisms that can be associated with hair loss [7], we were unable to find evidence of any microorganism. This is by no means indicative that an infectious agent can be discarded as causal agents. However, at least with the methods here used, we can establish that microorganisms commonly associated with alopecia were not present. Other microorganisms, such as some viruses, that have been associated with hair loss in pinnipeds [2,8] can damage hair indirectly due to their replication that leads to inflammation and epidermal lesions that can involve hair follicles. However, we did not find any evidence of skin lesions during our clinical examination of the fur seals that were handled. Future analyses could use molecular techniques to further examine the possibility that dermatophytes and other currently unknown infectious agents could be directly or indirectly related to the cases of alopecia in Guadalupe fur seals.

It is particular interest to point out that during our surveys we did not find evidence of alopecia in any of the fur seal pups or in sexually mature individuals. However, this is in no way indicative of the condition been exclusive to juvenile males. In the study site, around 84% of the fur seals are juveniles [18,22], making it more likely to have observed alopecia in this age group.

If Guadalupe fur seal alopecia is the reflection of a metabolic syndrome caused by biochemical alterations arising from nutritional deficiencies consequential to climatic anomalies, then the prevalence of alopecia would likely exhibit temporal variations, as occurs with other diseases [56]. It is also, of course, possible that the observed alopecia is related to an unidentified systemic condition that in turn is also sensitive to environmental factors [1,3]. Future long-term studies that take into account nutritional and immune biomarkers would help to ascertain this possibility. Given the increased frequency of climatic anomalies that have been recorded in the past decades [31], and the increased exposure of marine mammals to xenobiotics that can impact their metabolism and immune function [57], that further exacerbate the biological consequences of climatic anomalies, it is pressing to increase our knowledge of physiology and health of free living wildlife, particularly of species whose conservation is already endangered and that could be particularly susceptible to such changes.

## Acknowledgements

All procedures were conducted by approval of the Bioethics Committee and IACUC of the Autonomous University of Queretaro (Approval No. 59FCN2021). Sampling was conducted under permits SGPA/DGVS/00091/17 and SGPA/DGVS/01643/19 issued by the Dirección General de Vida Silvestre. We thank the Abalone Fishermen Cooperative (Cooperativa Nacional de Abuloneros) for their logistical support. Luis A. Soto-García, Cecilia Barragán-Vargas, Fabiola Guerrero de la Rosa, and Hiram Rosales Nanduca provided assistance during fieldwork. Field work was partly funded under a CONACYT Research Grant (A1-S-16417) awarded to K.A.W for research on pinnipeds.

